# Diverse ssRNA viruses associated with *Karenia brevis* harmful algal blooms in Southwest Florida

**DOI:** 10.1101/2025.01.06.631608

**Authors:** Shen Jean Lim, Alexandra Rogers, Karyna Rosario, Makenzie Kerr, Matt Garrett, Julie Koester, Katherine Hubbard, Mya Breitbart

## Abstract

Harmful algal blooms (HABs) caused by the dinoflagellate *Karenia brevis* frequently occur in the eastern Gulf of Mexico, where they negatively impact the environment, human health, and economy. Very little is known about viruses associated with *K. brevis* blooms, although viral infection of other HAB-forming phytoplankton species can play an important role in bloom dynamics. We used viral metagenomics to identify viruses in 11 pooled seawater samples collected from Southwest Florida, USA in 2021 during a severe, spatiotemporally dynamic *K. brevis* bloom. Assembled viral genomes were similar to published genomes from the order *Picornavirales*, family *Marnaviridae*, and genera *Sogarnavirus*, *Bacillarnavirus*, and *Marnarnavirus*. Viruses from these groups infect bloom-forming diatoms (*Chaetoceros* sp. and *Rhizosolenia setigera*) and the raphidophyte *Heterosigma akashiwo*. We also recovered unclassified *Riboviria* genomes related to a *Symbiodinium* positive-sense ssRNA virus sequenced from coral dinoflagellate symbionts. Reverse-transcriptase PCR assays were performed to monitor the occurrence of seven representative virus genomes in these 2021 samples and 43 seawater samples collected during a subsequent, typical bloom between November 2022 and May 2023. Over half of the samples contained multiple viruses, and at least one viral genome was detected in 44 of 54 samples, collected across seasons and years, highlighting the ubiquity of these viruses in this region. Alpha diversity was highest in the summer months and positively correlated with *K. brevis* cell counts. Multiple regression revealed month and the presence of unclassified *Riboviria* sequences most similar to dinoflagellate viruses as significant predictors of *K. brevis* cellular abundance.

**Importance:** Harmful algal blooms caused by the dinoflagellate *Karenia brevis* negatively impact the tourism, fisheries, and public health sectors. Anticipated impacts of climate change, nutrient pollution, and ocean acidification may sustain and/or exacerbate *K. brevis* blooms in the future, underscoring the need for proactive monitoring, communication, and mitigation strategies. This study represents a pioneering effort in monitoring viruses associated with *K. brevis* blooms. The findings lay the groundwork for studying the effects of environmental drivers on *K. brevis* blooms and their associated viruses, as well as for exploring the roles of viruses in bloom dynamics and potential applications of viruses as biocontrol agents for *K. brevis* blooms. Furthermore, the comparison of viral dynamics relative to local and regional bloom dynamics in this study helps inform future monitoring and modeling needs.

## Introduction

The dinoflagellate species *Karenia brevis* forms harmful agal blooms (HABs), commonly known as “red tides”, almost annually along the coasts and offshore waters of the eastern Gulf of Mexico (1, 2). Extensive *K. brevis* blooms cause water discoloration and, in some cases, hypoxia (3). *Karenia brevis* also produces brevetoxins that exhibit neurotoxic (4) or hemolytic (5) activity. These brevetoxins are ichthyotoxic and cause fish kills upon direct exposure (6). Brevetoxins and their metabolites can also accumulate in filter-feeding zooplankton and shellfish, where they are transferred up the food chain (7, 8). This trophic transfer contributes to fish, bird, and marine mammal mortalities, as well as Neurotoxic Shellfish Poisoning (NSP) in humans (9). *Karenia brevis* cells are readily lysed by high turbulence at the water surface or along the shore, releasing brevetoxins into the water (10). Released brevetoxins can be further aerosolized and transported onshore by winds, where they trigger respiratory irritation in animals (11, 12) and humans (9, 13). The negative aesthetic, environmental, and health effects of *K. brevis* blooms lead to significant economic losses across the tourism, fisheries, and public health sectors (14). These impacts are evident in the Gulf of Mexico, particularly on the west coast of Florida, which experiences the most frequent and persistent occurrences of *K. brevis* blooms (2).

Florida prioritizes strategies that aim to reduce the ecological, health, and economic consequences of *K. brevis* blooms (15). These include collaborative efforts for monitoring and forecasting blooms and related impacts, such as the FWC Historical HAB Database which compiles and hosts *K. brevis* cell abundance data from 1953 to the present (https://myfwc.com/research/redtide/monitoring/database/), specialized remote sensing techniques (16), and ocean circulation models (17). These data are integrated to inform shellfish harvesting management (18), forecasts published by NOAA’s National Centers for Coastal Ocean Science (https://coastalscience.noaa.gov/science-areas/habs/hab-forecasts/gulf-of-mexico/) and the University of South Florida (19), and related public outreach initiatives. Biological, chemical, and physical factors are thought to underlie variability in bloom severity and duration (6, 20, 21). However, accurately modeling complex bloom dynamics has proven challenging and is an ongoing process (22). It is critical to identify and/or parameterize ecological metrics and strategies that are relevant for growth and loss at varying scales. This is helpful for building complexity into forecasting models and hindcasting past events.

*Karenia brevis* blooms can endure for a few months to a few years (23) and can be localized to a single estuary, or span Florida’s Gulf and Atlantic coasts (2). A *K. brevis* bloom event can be broken down into stages, including initiation, growth, maintenance, decline, and termination (21). These stages can occur regionally to locally; for example, a single bloom may initiate only once but may pass through a certain area multiple times during a bloom event. In addition to physical and chemical influences, biotic mechanisms are thought to play important roles in various *K. brevis* bloom stages. These include interactions between *K. brevis* and grazers, parasites, other algal species, algacidal bacteria, and lytic viruses, as well as responses of *K. brevis* to other internal and external triggers, such as life history stage and photosynthesis (6, 20, 21). Understanding biological interactions between HAB species and microbes is crucial, as these interactions can critically influence bloom dynamics and biogeochemical cycling. Furthermore, climate change, with its cascading effects on water temperatures, ocean stratification, currents, and nutrient transport, is predicted to alter HAB interactions, dispersal, and frequency (24).

Virus-host interactions can shape HAB life cycles and serve important roles in bloom ecology (24). Factors such as viral lysis can result in bloom decline and may provide useful insights into bloom prevention and treatment approaches (2, 15). While both DNA and RNA viruses can infect HAB species (25), most research has focused on single-stranded RNA (ssRNA) viruses, which are prevalent in HAB-forming dinoflagellate, diatom, and raphidophyte species (26). To date, viruses associated with *K. brevis* blooms have not yet been sequenced. An unknown microorganism with lytic activity against *K. brevis* cultures was previously recovered from surface waters containing *K. brevis* blooms in Southwest Florida (27). These *K. brevis* cultures contained virus-like particles (VLPs) in their supernatants. However, lysing cells contained mostly bacteria and rarely VLPs, suggesting that *K. brevis* cell death may be caused by virus-bacteria interactions.

In this study, we aimed to identify and monitor RNA viruses in seawater from Southwest Florida associated with *K. brevis* blooms. Using viral metagenomics, we recovered near-complete viral genomes from surface water samples collected from 11 locations around Pinellas and Manatee counties in April, June, and July 2021, during a long-lasting and severe *K. brevis* bloom that began the prior year. We then designed reverse-transcriptase PCR (RT-PCR) primers for each representative viral genome to analyze their presence and ecology in the 2021 samples plus 43 additional seawater samples sampled from 39 locations around Southwest Florida during a subsequent *K. brevis* bloom event occurring between November 2022 and May 2023. Blooms of *K. brevis* typically start during fall and terminate in or before spring, and as in 2021, occasionally persist through summer. Herein, we use a bloom event to describe the bloom from initiation through termination, in recognition that blooms typically span multiple seasons and years.

## Methods

### Virome preparation

Seawater samples for virome sequencing were collected from four sites in April 2021, three sites in June 2021, and four sites in July 2021 (**Figure 1, Table 1, and Table S1**). From each site, 500 mL of surface water was collected in an acid-washed or autoclaved bottle and transported to the laboratory on ice. In the lab, each water sample was filtered through a 50 mm 0.45 μm Nalgene™ Rapid-Flow™ Sterile Disposable Filter Unit with PES membrane (Thermo Scientific™, Waltham, MA, USA). Each filter was placed in a sterile disposable petri dish and cut in half with a sterile razor blade. Half of the filter was homogenized for virome sequencing, while the other half was left in the petri dish, which was wrapped with aluminum foil and parafilm and stored at −80°C for RT-PCR. To ensure complete homogenization, the half of the filter used for virome sequencing was cut evenly into four sections. Each section was homogenized in a separate Zymo Research (ZR) BashingBead Lysis Tube (Irvine, CA, USA) containing 2 mm ceramic beads and 1 mL Suspension Medium (100 mM NaCl, 50 mM Tris-HCl (pH=7.5), and 8 mM MgSO_4_) for 1.5 minutes at maximum speed (5 meters/s) using a Fisherbrand^TM^ Bead Mill 4 Homogenizer (Fisher Scientific, Waltham, MA, USA). Each resulting homogenate was briefly centrifuged and 950 μL aliquots from each of the four homogenates per sample were pooled for the purification of virus-like particles (VLPs). Each pooled homogenate was centrifuged at 10,000 *g* for 10 minutes at 4°C and the supernatant was syringe-filtered through a 0.45-μm Sterivex filter (MilliporeSigma, Burlington, MA, USA). Half of the filtrate was treated with a nuclease mixture of 21 U of TURBO^TM^ DNase (Invitrogen^TM^, Waltham, MA, USA), 4.5 U of Baseline-ZERO^TM^ DNase (Epicentre, Paris, France), 112.5 U Benzonase® endonuclease (MilliporeSigma), and 400 U Ambion^TM^ RNase I in 1X Turbo^TM^ DNase Buffer (Invitrogen^TM^) for 1.5 hours at 37℃, followed by nuclease inactivation with 20 mM EDTA (pH=8.0) for identification of RNA viruses. The other half of the filtrate was treated similarly but without RNase I for future identification of DNA viruses.

**Figure 1.**
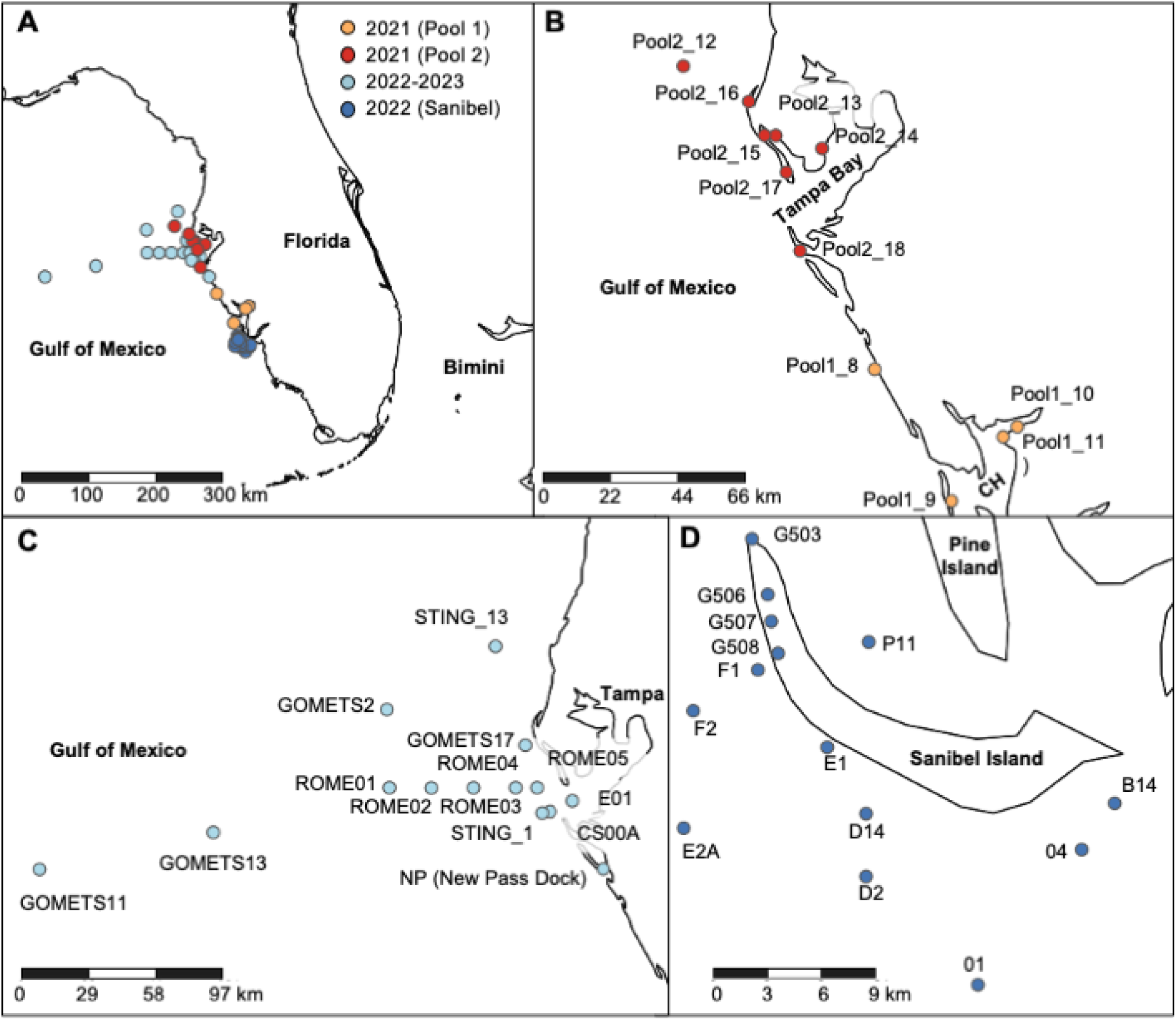
Map of locations of (a) all sampling sites, including (b) sites from the greater Charlotte Harbor (CH) and Tampa Bay regions where Pool 1 and Pool 2 samples were collected for virome sequencing in 2021; (c) other sites in, around, and offshore of Tampa Bay where samples were collected from 2022-2023; and (d) Sanibel Island sites where samples were collected in 2022. Seawater samples from all sites were collected for RT-PCR and their associated metadata are described in **Table 1** and **Table S1**.

**Table 1.** Metadata of samples analyzed in this study. Samples used for virome sequencing are highlighted in bold. Samples used for pooled libraries are indicated with the prefixes Pool1 and Pool2. Samples from Pool2 that were subsequently sequenced individually are indicated with *. *Karenia brevis* cell counts, available from the FWC HAB Monitoring Database (https://geodata.myfwc.com/datasets/myfwc::recent-harmful-algal-bloom-hab-events), were matched to the exact or closest (indicated with ^ in the last column) sample collection date, depth, latitude, and longitude. Metadata of samples used for *K. brevis* counts are provided in **Table S1**.

| Sample ID | Collection Date | Location | Latitude | Longitude | Depth (m) | *K. brevis* (cells/L) |
| --- | --- | --- | --- | --- | --- | --- |
| Pool1_8 | 4/6/2021 | Venice | 27.1091 | -82.4773 | 0.5 | 46,500^ |
| Pool1_9 | 4/6/2021 | Boca Grande | 26.7188 | -82.2488 | 0.5 | 17,000^ |
| Pool1_10 | 4/29/2021 | Laishley Park – boat ramp | 26.93939 | -82.052278 | 0.5 | 11,000 |
| Pool1_11 | 4/29/2021 | Ponce de Leon Park – boat ramp | 26.90927 | -82.09481 | 0.1 | 285,311 |
| Pool2_12* | 6/13/2021 | Gulf of Mexico – edge of offshore high Chl-a patch | 28.00568 | -83.04386 | 1 | 0^ |
| Pool2_13* | 6/17/2021 | War Veteran's Memorial Park | 27.80192 | -82.7746 | 0.5 | 1,298,692 |
| Pool2_14* | 6/30/2021 | Bayboro Harbor | 27.761897 | -82.6329 | 0.5 | 393,333^ |
| Pool2_15* | 7/25/2021 | Archibald Beach Park | 27.8009 | -82.8034 | 0.5 | 1,769,218 |
| Pool2_16* | 7/25/2021 | Indian Rocks Beach | 27.9006 | -82.8499 | 0.5 | 152,667 |
| Pool2_17* | 7/25/2021 | Pass-A-Grille Beach | 27.69141 | -82.7386 | 0.5 | 390,333 |
| Pool2_18* | 7/26/2021 | W of Cortez Beach (10th Street Pier) | 27.4595 | -82.6982 | 0.5 | 152,667 |
| NP1 | 11/7/2022 | New Pass Dock (Sarasota Bay) | 27.33375 | -82.57938 | 0.5 | 431,030 |
| NP2 | 11/14/2022 | New Pass Dock (Sarasota Bay) | 27.33375 | -82.57938 | 2.7 | 76,667 |
| NP3 | 5/8/2023 | New Pass Dock (Sarasota Bay) | 27.33375 | -82.57938 | 2 | 667 |
| ROME01_022423_surface | 2/24/2023 | 41 mi W of Bunces Pass | 27.64966 | -83.410366 | 0.5 | 0 |
| ROME01_032723_surface | 3/27/2023 | 41 mi W of Bunces Pass | 27.64966 | -83.410366 | 0.5 | 0 |
| ROME02_022423_surface | 2/24/2023 | 31 mi W of Bunces Pass | 27.64966 | -83.2474 | 0.5 | 0 |
| ROME02_032723_surface | 3/27/2023 | 31 mi W of Bunces Pass | 27.64966 | -83.2474 | 0.5 | 0 |
| ROME03_022423_surface | 2/24/2023 | 21 mi W of Bunces Pass | 27.64966 | -83.084434 | 0.5 | 1,333 |
| ROME03_032723_surface | 3/27/2023 | 21 mi W of Bunces Pass | 27.64966 | -83.084434 | 0.5 | 333 |
| ROME04_120522_surface | 12/5/2022 | 11 mi W of Bunces Pass | 27.64966 | -82.921468 | 0.5 | 4,752,180 |
| ROME04_022423_surface | 2/24/2023 | 11 mi W of Bunces Pass | 27.64966 | -82.921468 | 0.5 | 12,000 |
| ROME04_032723_surface | 3/27/2023 | 11 mi W of Bunces Pass | 27.64966 | -82.921468 | 0.5 | 41,333 |
| ROME05_022423_surface | 2/24/2023 | 5.8 mi W of Bunces Pass | 27.64966 | -82.84019 | 0.5 | 1,719,010 |
| ROME05_022423_bottom | 2/24/2023 | 5.8 mi W of Bunces Pass | 27.64966 | -82.84019 | 8.8 | 1,333 |
| ROME05_032723_surface | 3/27/2023 | 5.8 mi W of Bunces Pass | 27.64966 | -82.84019 | 0.5 | 94,000 |
| ROME05_032723_bottom | 3/27/2023 | 5.8 mi W of Bunces Pass | 27.64966 | -82.84019 | 8.8 | 11,333 |
| ROME05_042523_surface | 4/25/2023 | 5.8 mi W of Bunces Pass | 27.64966 | -82.84019 | 0.5 | 0 |
| ROME05_042523_bottom | 4/25/2023 | 5.8 mi W of Bunces Pass | 27.64966 | -82.84019 | 8.7 | 0 |
| Sanibel_01 | 11/17/2022 | 5.8 mi S of Tarpon Bay Road Beach | 26.33856 | -82.09553 | 0.5 | 576,810 |
| Sanibel_04 | 11/17/2022 | 2.4 mi SE of Tarpon Bay Road Beach | 26.40238 | -82.04633 | 0.5 | 1,462,680 |
| Sanibel_B14 | 11/17/2022 | 2.1 mi SW of Lighthouse Beach Park | 26.42421 | -82.03075 | 0.5 | 314,167 |
| Sanibel_D2 | 11/17/2022 | 4.5 mi S of Bowmans Beach | 26.3897 | -82.14855 | 0.5 | 75,000 |
| Sanibel_P11 | 11/17/2022 | 1.2 mi E of Chino Island (Pine Island Sound) | 26.500267 | -82.1473 | 0.5 | 24,667 |
| Sanibel_D14 | 11/17/2022 | 2.5 mi S of Bowmans Beach | 26.41931 | -82.14855 | 0.5 | 234,125 |
| Sanibel_E1 | 11/17/2022 | SW of Bowmans Beach | 26.45062 | -82.16691 | 0.5 | 11,333 |
| Sanibel_E2A | 11/17/2022 | 5.8 mi SW of Bowmans Beach | 26.41249 | -82.23468 | 0.5 | 392,551 |
| Sanibel_F1 | 11/17/2022 | 1 mi NW of Blind Pass | 26.48713 | -82.19962 | 0.5 | 104,055 |
| Sanibel_F2 | 11/17/2022 | 3.1 mi W of Blind Pass | 26.46784 | -82.23019 | 0.5 | 20,000 |
| Sanibel_G503 | 11/17/2022 | SW of Redfish Pass | 26.54911 | -82.20238 | 0.5 | 283,667 |
| Sanibel_G506 | 11/17/2022 | SW of Alison Hagerup Beach | 26.52273 | -82.19497 | 0.5 | 157,284 |
| Sanibel_G507 | 11/17/2022 | W of Captiva Island Yacht Club | 26.50999 | -82.19332 | 0.5 | 239,667 |
| Sanibel_G508 | 11/17/2022 | W of Osprey Way Drive | 26.49495 | -82.19012 | 0.5 | 196,905 |
| STING_1_1m | 2/20/2023 | STING Station 1 - CTD | 27.5515 | -82.8184 | 1 | 398,000^ |
| STING_1_8m | 2/20/2023 | STING Station 1 - Pump | 27.5515 | -82.8184 | 8 | 315,000^ |
| STING_13_2m | 2/26/2023 | STING Station 13 | 28.2006 | -83.0001 | 2 | 0^ |
| E01 Surface | 4/25/2023 | 1.8 mi S of Mullet Key (Lower Tampa Bay) | 27.5991 | -82.7 | 0.5 | 0 |
| E01 Bottom | 4/25/2023 | 1.8 mi S of Mullet Key (Lower Tampa Bay) | 27.5991 | -82.7 | 5.9 | 0 |
| CS00A Surface | 4/25/2023 | 2.3 mi SW of Egmont Key | 27.55661 | -82.78781 | 0.5 | 0 |
| CS00A Bottom | 4/25/2023 | 2.3 mi SW of Egmont Key | 27.55661 | -82.78781 | 7.2 | 0 |
| GOMETS2 | 5/8/2023 | GOMETS Station 2 | 27.95464 | -83.42188 | 1 | 0^ |
| GOMETS11 | 5/12/2023 | GOMETS Station 11 | 27.3325 | -84.7740 | 1 | 0^ |
| GOMETS13 | 5/12/2023 | GOMETS Station 13 | 27.4762 | -84.0963 | 1 | 0^ |
| GOMETS17 | 5/13/2023 | GOMETS Station 17 | 27.8153 | -82.8835 | 1 | 0^ |

NGS library preparation was performed using methods described in Rosario *et al.* (2022) (28). Briefly, 50 µL of RNA was extracted from each purified RNA virus fraction using Qiagen’s RNeasy Mini kit (Valencia, CA, USA) with on-column DNase digestion and quantified using the Qubit^TM^ RNA high sensitivity (HS) assay. Two RNA samples pooled from seawater samples collected in April 2021 and June-July 2021 (**Table 1**) were initially processed for sequencing. Subsequently, seven individual RNA samples from both pools were processed to improve the coverage of viral sequences for genome assembly. From each sample, cDNA was synthesized from 8 μL RNA and 50 ng random hexamers using the SuperScript^TM^ IV First-Strand Synthesis System (Invitrogen^TM^). Second strand synthesis was performed on each cDNA product using New England Biolabs’ DNA Polymerase I, Large (Klenow) Fragment (Ipswich, MA, USA). Double-stranded cDNA samples were purified with ZR’s DNA Clean & Concentrator-25 kit and fragmented to 300 bp at the University of South Florida’s Genomics Program Sequencing core (Tampa, FL, USA) using the Covaris® M220 Focused-ultrasonicator (Woburn, MA, USA). NGS libraries were prepared from fragmented cDNA samples using the xGen™ ssDNA & Low-Input DNA Library Preparation Kit (Integrated DNA Technologies, Coralville, IA, USA) following the manufacturer’s protocol for DNA concentrations <1 ng/μL. All libraries were paired-end sequenced (2 x 150 bp) by Azenta Life Sciences (Burlington, MA, USA) on an Illumina HiSeq platform.

### Virome assembly and sequence analysis

Sequencing reads were trimmed using the default parameters of Trimmomatic v0.39 (29) with a custom read head crop of 10 bases, as described in Rosario *et al.* (2022) (28). Read quality pre- and post-trimming, including the presence of adapter sequences, was evaluated using FastQC v0.11.8 (https://www.bioinformatics.babraham.ac.uk/projects/fastqc). To obtain near-complete or complete draft viral genomes, several assembly approaches were used. Trimmed reads from each pooled cDNA library (April 2021 and June-July 2021) were initially assembled using the pipeline recommended for PCR-amplified viral metagenomes (30) which included read deduplication using BBtool’s clumpify.sh (dedupe subs=0 passes=2; https://sourceforge.net/projects/bbmap), followed by assembly using single-cell SPAdes (31). Trimmed reads from each individual and pooled cDNA library were also individually assembled using the default options of the newer rnaviralSPAdes assembler specific for RNA viruses (32) integrated in SPAdes v3.15.5 (31). Additionally, reads from all sequenced libraries were also combined, deduplicated using BBtool, and co-assembled using rnaviralSPAdes (32).

RNA viral contigs were identified from each assembly using VirBot (33). Sequences of RNA viral contigs ≥1,000 nt were compared with sequences from NCBI’s nucleotide (nt) and non-redundant (nr) protein collections (34) using the megablast and/or blastx programs on NCBI’s Basic Local Alignment Research Tool (BLAST®) server (35) to identify genome relatives. Contigs homologous to RNA viruses associated with eukaryotic phytoplankton were assessed for quality and completeness using the default parameters of CheckV v1.0.1 (36). Draft genomes with ≥80% completeness (**Table 2**) were retained for assembly correction, genome annotation, and primer design. For assembly correction, trimmed reads from each library were first mapped to each contig using the default options of bowtie2 v2.2.5 (37). Each resulting SAM (Sequence Alignment Map) file was converted to a sorted and indexed BAM (Binary Alignment Map) using samtools v1.9 (38). The sequence and BAM alignments of each contig were provided as input to Pilon v1.24 (39) to automatically fix nucleotide differences, insertions/deletions, gaps, and local assemblies in the draft genome, using the default parameters. For each base in the input genome, Pilon (39) evaluates whether the base call is supported by a majority of evidence from read alignments, which are weighted by base and mapping quality. Open reading frames (ORFs) in each corrected genome were identified using NCBI’s Open Reading Frame Finder (ORF Finder) web tool (https://www.ncbi.nlm.nih.gov/orffinder). For gene/protein annotation, genome sequences were compared against annotated RNA helicase, RNA-dependent RNA polymerase (RdRp), and viral protein (VP) sequences in related genomes using the tblastn program on NCBI’s BLAST® server (35). To annotate the conserved structural protein domains VP1-VP3, reference protein sequences from *Heterosigma akashiwo* RNA virus (HaRNAV) were used (40). Genome organization was visualized using the ggplot2 v3.5.1 (41) R package with the gggenes v0.5.1 extension (https://doi.org/10.32614/CRAN.package.gggenes).

**Table 2.** Features of ssRNA viral genomes assembled in this study. Representative genomes for each viral taxon are indicated with asterisks (*) in the first column.

| Genome ID | NCBI Accession | Library | Assembly method | Genome length (nt) | Genome completeness (%) | Taxonomy | Host Species (type) |
| --- | --- | --- | --- | --- | --- | --- | --- |
| Riboviria1_1* | PP297088 | All pooled | Co-assembly (rnaviralSPAdes) | 6,078 | 100 | *Viruses;Riboviria* | Unknown (dinoflagellate) |
| Riboviria1_2 | PP297086 | Pool2 | rnaviralSPAdes | 5,901 | 100 |  |  |
| Riboviria1_3 | PP297087 | Pool2 | single-cell SPAdes | 6,058 | 100 |  |  |
| Riboviria_2* | PP297085 | All pooled | Co-assembly (rnaviralSPAdes) | 4,158 | 91.3 | *Viruses;Riboviria* | Unknown (dinoflagellate) |
| Sogarnavirus1_1* | PP297083 | Pool2 | single-cell SPAdes | 9,790 | 100 | *Viruses;Riboviria;Orthornavirae;Pisuviricota;Pisoniviricetes;Picornavirales;Marnaviridae;Sogarnavirus* | Unknown (diatom) |
| Sogarnavirus1_2 | PP297084 | All pooled | Co-assembly (rnaviralSPAdes) | 7,225 | 80.35 |  |  |
| Sogarnavirus2_1* | PP297079 | All pooled | Co-assembly (rnaviralSPAdes) | 9,652 | 100 | *Viruses;Riboviria;Orthornavirae;Pisuviricota;Pisoniviricetes;Picornavirales;Marnaviridae;Sogarnavirus;Chaetarnavirus 2* | *Chaetoceros* sp. (diatom) |
| Sogarnavirus2_2 | PP297078 | Pool2_14 | rnaviralSPAdes | 9,574 | 100 |  |  |
| Sogarnavirus2_3 | PP297080 | Pool2 | rnaviralSPAdes | 9,575 | 100 |  |  |
| Sogarnavirus3* | PQ118637 | All pooled | Co-assembly (rnaviralSPAdes) | 8,488 | 92.96 | *Viruses;Riboviria;Orthornavirae;Pisuviricota;Pisoniviricetes;Picornavirales;Marnaviridae;Sogarnavirus;Chaetarnavirus 2* | *Chaetoceros* sp. (diatom) |
| Bacillarnavirus1* | PP297082 | All pooled | Co-assembly (rnaviralSPAdes) | 8,983 | 96.82 | *Viruses;Riboviria;Orthornavirae;Pisuviricota;Pisoniviricetes;Picornavirales;Marnaviridae;Bacillarnavirus;Rhizosolenia setigera RNA virus 01* | *Rhizosolenia setigera* (diatom) |
| Marnavirus1* | PP297081 | All pooled | Co-assembly (rnaviralSPAdes) | 7,710 | 88.66 | *Viruses;Riboviria;Orthornavirae;Pisuviricota;Pisoniviricetes;Picornavirales;Marnaviridae;Marnavirus* | Unknown (raphidophyte) |

Nucleotide sequences of all newly assembled genomes were aligned with sequences of their genome relatives that were identified by BLAST using the L-INS-I and --adjustdirectionaccurately options on the MAFFT v7 server (42). Amino acid sequences of the RdRp and capsid protein were also extracted from the annotation data of each viral genome for Multiple Sequence Alignment (MSA). The MSA template used for RdRp sequences was a previously published MSA (43), spanning eight conserved domains (44), that was used by the International Committee on Taxonomy of Viruses (ICTV) for *Marnaviridae* genus/species demarcation (45). This RdRp reference MSA was reconstructed using the MAFFT v7 server (42), followed by alignment editing using Unipro UGENE v48.1 (46). The MSA template used for capsid protein sequences was downloaded from the Pfam database (47) (Pfam entry: PF08762). This protein family was identified using the InterProScan online tool (48), using capsid protein sequences assembled in this study as input. The Pfam capsid protein MSA, which contained 204 sequences, was pruned to retain 83 sequences with stable phylogenetic positions that are consistent with the phylogeny reported in (43) and in the ICTV Report on *Marnaviridae* (45). RdRp and capsid protein sequences assembled in this study were added to their respective reference alignments using the --add, --keeplength, and L-INS-I options on the MAFFT v7 server (42). Pairwise sequence identities were computed from each MSA, with alignment gaps included, using the “Similarity” algorithm of the “Generate distance matrix” function in Unipro UGENE v48.1 (46). Phylogenetic analysis was performed on the RdRp (118 aa) and capsid protein (746 aa) MSA according to the methods described in (43). PhyML 3.0 (49), as implemented on the NGPhylogeny.fr server (50), was used to generate a maximum likelihood tree from each MSA using the default LG+I+G+F amino acid model and default random seed of 123456. For tree topology search, the best of NNI (Nearest Neighbor Interchange) or SPR (Subtree Pruning and Regraphing) was selected. Branch support was determined using the Shimodaira-Hasegawa (SH)-like statistical test (51). All trees were annotated using FigTree v1.4.4 (http://tree.bio.ed.ac.uk/software/Figtree).

For read coverage analysis, the longest genome of each viral taxon (**Table 2**) was selected as the representative genome and used to build a Bowtie index using bowtie2 v2.5.0 (37) and a bin collection in anvi’o v8 (52). Trimmed reads from each library were mapped to the Bowtie2 index and converted into a sorted and indexed BAM file using bowtie2 (37) and samtools v1.16.1 (38). All generated BAM files were imported into anvi’o (52) using the anvi-profile command and merged into a single profile using the anvi-merge command. To minimize non-specific mapping, the mean Q2Q3 coverage, calculated as the average depth of coverage for nucleotide positions that fall between the 2^nd^ and 3^rd^ quartiles of the coverage distribution for each genome, across each representative genome were calculated from the merged profile using the anvi-summarize command.

### Primer design

RT-PCR primers were designed to amplify conserved genes in the representative genomes of each viral taxon using the modified Primer3 software implemented in the Geneious Prime 2023 full release (53) or the PrimerQuest^TM^ tool in the Integrated DNA Technologies (IDT, Skokie, IL, USA) web server, with the following parameters: Tm: 58-60 °C, GC content: 50-60%, and primer size: 15-30 bp. Candidate primer sequences (**Table S2**) were evaluated for genome specificity via sequence searches using Unipro UGENE v48.1 (46) against all genomes assembled in this study, which include target and non-target genomes, as well as sequence searches against all organisms in NCBI’s nr nucleotide collection (34) using the default parameters of Primer-BLAST (54). Newly designed primer pairs were tested and optimized on pooled cDNA from the 11 seawater samples used for virome sequencing. A gradient PCR reaction was performed to determine the optimal annealing temperature of each primer pair. The reaction contained 0.48 μM of each primer, 2 μL cDNA template, 1 μL GC enhancer, and 1X AmpliTaq Gold™ 360 Master Mix (Applied Biosystems™, Waltham, MA, USA) in a 25 μL reaction volume. Gradient PCR was performed under the following conditions: Initial denaturation at 95°C for 10 minutes, 40 cycles of denaturation at 95°C for 30 seconds, annealing at 53-59.9°C for 30 seconds, extension at 72°C for 1 minute, followed by elongation at 72°C for 10 minutes and cooling at 11°C. PCR products were visualized following gel electrophoresis on a 2% (wt/vol) agarose gel stained with ethidium bromide. For each primer set, the annealing temperature that produced the brightest band with no nonspecific amplification was chosen for subsequent RT-PCR assays. PCR products were purified using the Zymoclean Gel DNA Recovery Kit, quantified using the Qubit^TM^ DNA high sensitivity (HS) assay (Invitrogen™), and Sanger sequenced bidirectionally by TACGen (Richmond, CA, USA) or Eurofin Genomics (Louisville, KY, USA).

### RT-PCR

RT-PCR assays using each newly designed primer pair were performed on the 11 samples used for virome sequencing, 43 additional seawater samples collected during a subsequent *K. brevis* bloom event, and 62 samples from cell cultures of *Karenia* spp. (**Table 1** and **Table S3**). The 43 seawater samples were collected by the Fish and Wildlife Research Institute (FWRI) from around southwest Florida between 2022 and 2023 (**Table 1**). The volume of collected seawater samples ranged from 500 mL to 2 L, and each sample was filtered through a 50 mm, 0.45 μm filter as described above. The 62 cell culture samples were obtained from 17 strains of *Karenia* spp. grown in three independent flasks at FWRI (**Table S3**); two replicates were sampled in mid-exponential phase and the third was harvested in stationary phase. *Karenia brevis* strains CCFWC 242, 254, 257, 258 (only two replicates available), 261, 267, 121, 123, 124, 125, 126, 1010, 1012, 1013, 1014, 1016, and 1021, *K. mikimotoi* strain CCFWC 67, *K. papilionacea* strain CCFWC 1020, *K. umbella* strain 1019 B2, and media blanks were individually filtered through 25 mm, 0.45 μm Durapore® PVDF membrane filters (MilliporeSigma). The volume of culture filtered ranged from 9 mL to 33 mL. From each filtered field-collected or culture sample, a quarter of each filter was cut out, homogenized, and used for total RNA extraction, quantification, cDNA synthesis, and RT-PCR (**Table S2**) using the methods described above.

### Viral diversity analysis

Location data for field-collected samples was used to create maps with SimpleMappr (https://www.simplemappr.net). *Karenia brevis* counts were downloaded from FWRI’s harmful algal bloom (HAB) database at https://geodata.myfwc.com/datasets/myfwc::recent-harmful-algal-bloom-hab-events. Cell counts for each field-collected sample were matched to the exact or closest sample collection date, depth, latitude, and longitude. RT-PCR results from all field-collected samples were combined to generate a presence/absence matrix for diversity analysis, where presence=1 and absence=0. The RT-PCR data matrix and sample metadata were separately imported into R v4.0.2. Diversity analysis was performed using the ampvis2 v2.7.34 R package (55), which converted the RT-PCR data into an Operational Taxonomic Unit (OTU) table, where each OTU indicates a representative genome from each viral taxon (**Table 2**). This OTU table was linked to the sample metadata and stored as an ampvis2 object. Alpha diversity was computed with the amp_alphadiv function using the observed OTU measure. Shapiro-Wilk tests, implemented in R, rejected the null hypothesis (*p*<0.05) that alpha diversity and numerical environmental variables (latitude, longitude, depth, and *K. brevis* cell counts) were normally distributed. Therefore, non-parametric statistical tests were used for downstream analyses with a statistical significance threshold of *p*<0.05. Alpha diversity between sample groups, clustered by year, season, month, or *K. brevis* cell count category, was compared using Kruskal-Wallis rank sum tests (56) implemented in R. *Karenia brevis* cell count category was defined according to FWRI’s criteria (https://www.flickr.com/photos/myfwc/sets/72157635398013168) of not present/background (0-1,000 *K. brevis* cells/L); very low (>1,000-10,000 *K. brevis* cells/L); low (>10,000-100,000 *K. brevis* cells/L); medium (>100,000-1,000,000 *K. brevis* cells/L); and high (>1,000,000 *K. brevis* cells/L). Correlations between alpha diversity and numerical environmental variables were evaluated using the Kendall rank correlation test implemented in the cor.test function in R. Kendall rank correlation was recommended for small, non-parametric datasets with many repeated ranks (57). Alpha diversity results were visualized using the ggplot2 v3.5.1 (41) R package with the ggbeeswarm v0.7.2 extension (https://doi.org/10.32614/CRAN.package.ggbeeswarm). Beta diversity was computed and visualized with ampvis2’s amp_ordinate function using the Principal Components Analysis (PCA) method with presence/absence transformation. Environmental factors were fitted onto the resulting PCA plot using the envfit function in the VEGAN v2.6.4 R package (58), which reports a goodness of fit R^2^ value and *p*-value for each variable.

### Multiple regression

Multiple regression models were fitted to analyze associations between predictor variables (environmental factors) and the response variable alpha diversity, as well as predictor variables (environmental factors and virus presence/absence) and the response variable *K. brevis* cell counts. All models were initially created using the lmp function of the lmperm v2.1.0 R package (https://doi.org/10.32614/CRAN.package.lmPerm), which uses permutation tests instead of the normal distribution to calculate *p*-values. Each model was optimized with a stepwise algorithm that adds and removes predictor variables from the model until it reaches the lowest possible Aikake Information Criterion (AIC) value (59), using the stepAIC function of the Modern Applied Statistics with S (MASS v7.3.60) R package (60). Multicollinearity between predictor variables in each optimized model was detected using a generalized variance inflation factor (GVIF) threshold of ≥4, which corresponds to GVIF ^(1/(2xdf))^ ≥2. GVIF values were calculated using the vif function in the MASS package (60), with the type argument set to “predictor”. Highly correlated variables, if any, were removed incrementally from the initial model and the model was re-optimized with stepAIC. This process was repeated until a final model with no multicollinearity was generated. Statistically significant predictors in each final model were identified using Analysis of Variance (ANOVA) tests implemented in R.

### Data availability

Sequenced reads and viral genomes were deposited in NCBI under the BioProject accession number PRJNA1149755. Viral genomes were assigned the GenBank accession numbers PP297078-PP297088 and PQ118637.

## Results

### Viral genomes

Sampling occurred during an unusually long bloom event from December 2020 to November 2021 (61), and at the time of year when most blooms have subsided (FWC Historical HAB Database). We sequenced the viromes of 11 surface seawater samples collected in Pinellas and Manatee counties in April (n=4), June (n=3), and July 2021 (n=4) (**Figure 1, Table 1**, and **Table S1**). Two pooled libraries comprising samples collected from the greater Charlotte Harbor region in April 2021 (Pool 1) and the greater Tampa Bay region in June-July 2021 (Pool 2) were initially sequenced and analyzed. Sequences related to ssRNA viruses infecting eukaryotic phytoplankton were only detected in Pool 2. To improve the coverage of viral sequences for genome assembly, viromes from each of the seven samples in Pool 2 were also individually sequenced (**Table 1 and Figure 1**). From these pooled and individually sequenced libraries, we assembled 12 viral genomes (**Table 2**) with ≥80% completeness using single-cell SPAdes (31) or rnaviralSPAdes (32).

Among the 12 assembled genomes, four genomes were most similar to viruses infecting dinoflagellates that remain unclassified within the *Riboviria*, including Riboviria1_1, Riboviria1_2, Riboviria1_3 and Riboviria2. (**Table 2**). The four genomes contained two open reading frames (ORFs) encoding the RNA-dependent RNA polymerase (RdRp) and the major capsid protein (MCP; **Figure 2a**). Based on CheckV’s (36) estimates, genomes Riboviria1_1, Riboviria1_2, and Riboviria1_3 were complete with an average genome size of 6,012±97 nt, while genome Riboviria2 was ∼91% complete and 4,158 nt in length (**Table 2**). Genomes Riboviria1_1, Riboviria1_2, and Riboviria1_3 shared 99±1% pairwise nucleotide identity, 97±2% RdRp sequence identity, and 100% capsid protein sequence identity with each other, but <60% pairwise nucleotide identity, <35% RdRp protein sequence identity, and 77% capsid protein identity to the genome Riboviria2 (**Data Set S1**). Based on their RdRp (**Figure 2b**) and capsid protein (**Figure S1**) phylogeny, the genomes Riboviria1_1, Riboviria1_2, and Riboviria1_3 were related to *Symbiodinium +ssRNA virus TR74740* (NCBI accessions: KX538960 and KX787934) (62), with which they shared 43%±6% RdRp sequence identity, but 80±1% capsid protein sequence identity (**Data Set S1**). The *Riboviria* genomes assembled in this study shared 22±3% RdRp sequence identity and 45±0% capsid protein sequence identity to the genome of *Heterocapsa circularisquama RNA virus 01* (HcRNAV; NCBI accession: NC_007518) that infects the HAB-forming dinoflagellate species *Heterocapsa circularisquama* (63).

**Figure 2.**
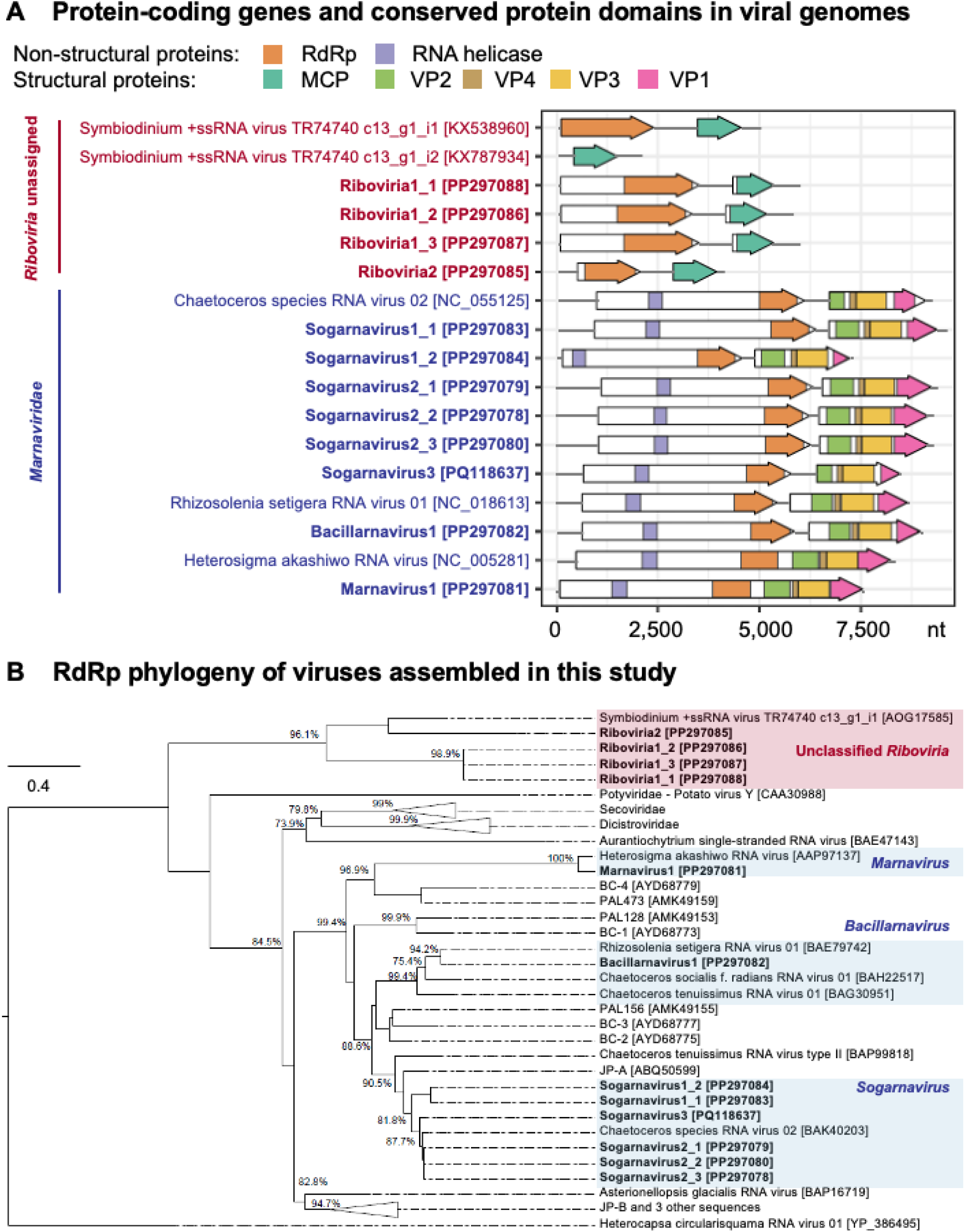
(a) Organization of open reading frames (ORFs; arrows) and their protein products (colored arrows) or conserved protein domains (colored subregions within white arrows) in genomes assembled in this study (bold text), in relation to related genomes from the realm unclassified *Riboviria* (red text) and the family *Marnaviridae* (blue text); (b) Maximum likelihood tree of RdRp amino acid sequences from viral genomes assembled in this study (bold text), in relation to sequences from related *Picornavirales* and unclassified *Riboviria* genomes. Sequences assigned to unclassified *Riboviria* are boxed in red, while sequences assigned to the order *Picornavirales*, family *Marnaviridae*, and genera *Marnavirus*, *Bacillarnavirus*, and *Sogarnavirus* are boxed in blue. The RdRp sequence of *Heterocapsa circularisquama RNA virus 01*, from the order *Sobelivirales*, is used as the outgroup. Node labels indicate SH-like branch support values >70%. The maximum-likelihood scale bar indicates the average number of substitutions per site. NCBI accession numbers for sequences in (a) and (b) are indicated in square brackets, where possible, and listed in **Data Set S1**.

The remaining eight genomes assembled in this study were most similar to phytoplankton-infecting viruses in the order *Picornavirales* and family *Marnaviridae* (**Table 2**). Genera in this family are defined by International Committee on Taxonomy of Viruses (ICTV) based on phylogenetic relationships of their RdRp protein sequences, while species are demarcated by a 90% RdRp sequence identity and 75% capsid protein sequence identity threshold (45). Our phylogenetic analysis assigned six assembled genomes to the genus *Sogarnavirus* and one genome each to the genera *Bacillarnavirus* and *Marnavirus* (**Figure 2b**).

The sogarnaviruses are represented by the complete 9,790-nt genome, Sogarnavirus1_1, and partial (∼80%) 7,225-nt genome, Sogarnavirus1_2. Both genomes shared 78% pairwise nucleotide identity, 99% RdRp sequence identity, and 88% capsid sequence identity with each other (**Table 2** and **Data Set S1**). Neither genome could be conclusively assigned to a species, because they shared 77±5% capsid protein sequence identity but <90% RdRp sequence identity with the *Chaetoceros species RNA virus 02* (CspRNAV02) genome (NCBI accession: NC_055125; unpublished), which is their closest relative (**Data Set S1**). The remaining sogarnavirus genomes, Sogarnavirus2_1, Sogarnavirus2_2, Sogarnavirus2_3, and Sogarnavirus3, were assigned to the species *Chaetarnavirus 2*, based on their 99±1% RdRp sequence identity and 85±3% capsid protein sequence identity to the CspRNAV02 genome. The complete genomes Sogarnavirus2_1, Sogarnavirus2_2, and Sogarnavirus2_3 had an average size of 9,600±45 nt (**Table 2**). They shared identical RdRp and capsid protein sequences and 99±0.6% pairwise nucleotide identity with each other, as well as 97% RdRp sequence identity, 79% capsid protein sequence identity, and 79±0.6% pairwise nucleotide identity with sequences in the Sogarnavirus3 genome, which was 8,488 nt long and ∼93% complete (**Table 2** and **Data Set S1**).

The genome assigned to the *Bacillarnavirus* genus, Bacillarnavirus1, was 8,983 nt long and ∼97% complete (**Table 2**). This genome was assigned to the species *Rhizosolenia setigera RNA virus 01*, based on its 96% RdRp sequence identity and 83% capsid protein sequence identity to the *Rhizosolenia setigera RNA virus 01* (RsRNAV01) reference genome (NCBI accession: NC_018613) (64). All genomes from the genera *Sogarnavirus* and *Bacillarnavirus* have di-cistronic genome organizations (**Figure 2a**), where the first ORF encodes non-structural proteins with the RNA helicase and RdRp domains, while the second ORF encodes the structural proteins VP2, VP4, VP3 and VP1 conserved in the family *Marnaviridae* (45).

Marnavirus1, the only genome representing the *Marnavirus* genus, was 7,710 nt long and ∼89% complete (**Table 2**). The species of this genome was unassigned, because its RdRp and capsid protein domain sequences were 86% and 94% identical, respectively, to the *Heterosigma akashiwo RNA virus* (HaRNAV) reference genome (NCBI accession: NC_005281) (40). Similar to the HaRNAV genome, genome Marnavirus1 has a mono-cistronic genome organization and encodes a single polyprotein with RNA helicase, RdRp, VP2, VP4, VP3, and VP1 protein sequence signatures (**Figure 2a**).

### Read coverage

Most viral genomes assembled in this study were only detected by sequencing in library Pool2, which included samples collected from the greater Tampa Bay region during June-July 2021 (**Table 3**). The only exception is the representative genome Sogarnavirus1_1, which was detected in both library Pool1 containing samples from the greater Charlotte Harbor region collected during April 2021, as well as library Pool2 (**Table 3**). Among all individually sequenced libraries from Pool2, the representative genomes Riboviria1_1, Riboviria2, Sogarnavirus1_1, Sogarnavirus3, and Bacillarnavirus1 had the highest coverage in library Pool2_13 (**Table 3**). This library was sequenced from a surface water sample collected at Boca Ciega Bay on June 2021 with ∼1.3 million *K. brevis* cells/L (**Table 1**). The highest coverage for the representative genomes Sogarnavirus2_1 and Marnavirus1 was in library Pool2_14 (**Table 3**). This library was sequenced from a surface water sample collected from Bayboro Harbor on June 2021 with ∼0.4 million *K. brevis* cells/L (**Table 1**).

**Table 3.** Average depth of read coverage of representative genomes from each viral taxon. To minimize non-specific mapping, only nucleotide positions with coverage values between the 2^nd^ and 3^rd^ quartiles (Q2Q3) of the coverage distribution for each genome were included. Coverage values of 0 are shaded in grey.

| Mean Q2Q3 Coverage | Pool1 | Pool2 | Pool2_12 | Pool2_13 | Pool2_14 | Pool2_15 | Pool2_16 | Pool2_17 | Pool2_18 |
| --- | --- | --- | --- | --- | --- | --- | --- | --- | --- |
| Riboviria1_1 | 0 | 63 | 0 | 125 | 0 | 0 | 0 | 0 | 0 |
| Riboviria2 | 0 | 22 | 0 | 38 | 0 | 1 | 0 | 0 | 0 |
| Sogarnavirus1_1 | 34 | 140 | 0 | 239 | 0 | 0 | 0 | 0 | 0 |
| Sogarnavirus2_1 | 0 | 223 | 0 | 0 | 1641 | 0 | 0 | 0 | 0 |
| Sogarnavirus3 | 0 | 23 | 0 | 36 | 14 | 0 | 0 | 0 | 0 |
| Bacillarnavirus1 | 0 | 25 | 0 | 38 | 0 | 0 | 0 | 0 | 0 |
| Marnavirus1 | 0 | 11 | 0 | 0 | 81 | 0 | 0 | 0 | 0 |

### RT-PCR assay design

Specific primer pairs (**Table S2**) were designed to develop reverse-transcriptase PCR (RT-PCR) assays for each representative viral genome (**Table 2**). All newly designed primers were initially tested on the 11 samples from Pool1 and Pool2 used for virome sequencing. Four of the seven primer pairs yielded PCR products from at least one Pool1 sample, while all seven primer pairs yielded PCR products from at least one Pool2 sample (**Table 4**). Sequences from the genomes Riboviria2, Bacillarnavirus1, and Marnavirus1 were not amplified in any Pool1 samples (**Table 4**). Overall, these primers showed higher sensitivity in detecting target sequences compared to virome sequencing. All viral genomes sequenced in the virome libraries (**Table 3**) were also successfully amplified by their genome-specific primers (**Table 4**). Furthermore, RT-PCR detected viral sequences in Pool1 and Pool2 samples that were not detected previously by virome sequencing (**Table 3** and **Table 4**).

**Table 4.** Presence (+) or absence (-) of sequence fragments from each genome in Pool1 and Pool2 samples used for virome sequencing, as determined by RT-PCR. For Pool2 samples, ‘++’ denotes the presence of a RT-PCR product and its corresponding sequence (with >0 mean Q2Q3 coverage) in the virome library (Table 2). Pool1 samples were not individually sequenced using viromics and had no sample-specific coverage information.

| Genome | Pool1_8 | Pool1_9 | Pool1_10 | Pool1_11 | Pool2_12 | Pool2_13 | Pool2_14 | Pool2_15 | Pool2_16 | Pool2_17 | Pool2_18 |
| --- | --- | --- | --- | --- | --- | --- | --- | --- | --- | --- | --- |
| Riboviria1_1/1_2/1_3 | + | - | - | - | + | ++ | - | - | + | + | + |
| Riboviria2 | - | - | - | - | - | ++ | - | ++ | + | + | - |
| Sogarnavirus1_1/1_2 | - | - | + | + | - | ++ | + | + | + | + | + |
| Sogarnavirus2_1/2_2/2_3 | + | + | - | - | - | - | ++ | + | + | - | + |
| Sogarnavirus3 | + | - | + | - | - | ++ | ++ | + | - | + | + |
| Bacillarnavirus1 | - | - | - | - | - | ++ | + | - | - | - | - |
| Marnavirus1 | - | - | - | - | - | - | ++ | - | - | - | - |

### Virus detection

Using these primers, RT-PCR assays were performed to screen for the presence of viral genomes in seawater samples collected during a more typical *K. brevis* bloom event documented from October 2022 to May 2023 (61). The screening included 43 additional samples collected at varying depths from 28 locations around Southwest Florida between November 2022 and May 2023 (**Table 1**). For viral diversity analysis, presence/absence data from these 43 samples were combined with data from the 11 samples used for virome sequencing. At least one viral genome was detected in all 11 samples collected in 2021, 15 of 16 samples collected in 2022, and 18 of 27 samples collected in 2023 (**Table S4**). Viruses were detected mainly in nearshore, surface (<3 m depth) water samples (**Figure 3** and **Table S4**). *Sogarnavirus* sp. and *Chaetarnavirus 2*, both from the genus *Sogarnavirus*, had the broadest distribution compared to other ssRNA viruses analyzed in this study. Sequence fragments from these genomes were amplified in 37 near-shore and open ocean samples collected from depths between 0.1 m to 8.8 m (**Figure 3d-f** and **Table S4**). Among viral genomes assigned to unclassified *Riboviria*, the viral taxon represented by Riboviria1 genomes was more widely distributed compared to the viral taxon represented by genome Riboviria2. Conserved sequence fragments from the Riboviria1 genomes were amplified from 22 samples, spanning the north and south of the Southwest Florida region, collected from varying depths (0.5 m-2.7 m) and dates (2021-2023) (**Figure 3a** and **Table S4**). In contrast, sequence fragments from the genome Riboviria2 were amplified from only four samples collected at surface (0.5 m) depth around St. Petersburg between June-July 2021 (**Figure 3b** and **Table S4**). Sequence fragments from the genome Bacillarnavirus1 was detected in 11 seawater samples, including two surface samples (0.5 m depth) collected from St. Petersburg (War Veteran’s Memorial Park and Bayboro Harbor) in June 2021, two samples collected at 0.5 m and 2.7 m depths from New Pass Dock in November 2022, and seven surface samples (0.5 m depth) collected from Sanibel Island in November 2022 (**Figure 3c** and **Table S4**). *Marnavirus* sp. was the least prevalent in the seawater samples, since sequence fragments from the genome Marnavirus1 were only amplified from two samples, including Pool2_14 from Bayboro Harbor and sample 01 from Sanibel Island (**Figure 3g**).

**Figure 3.**
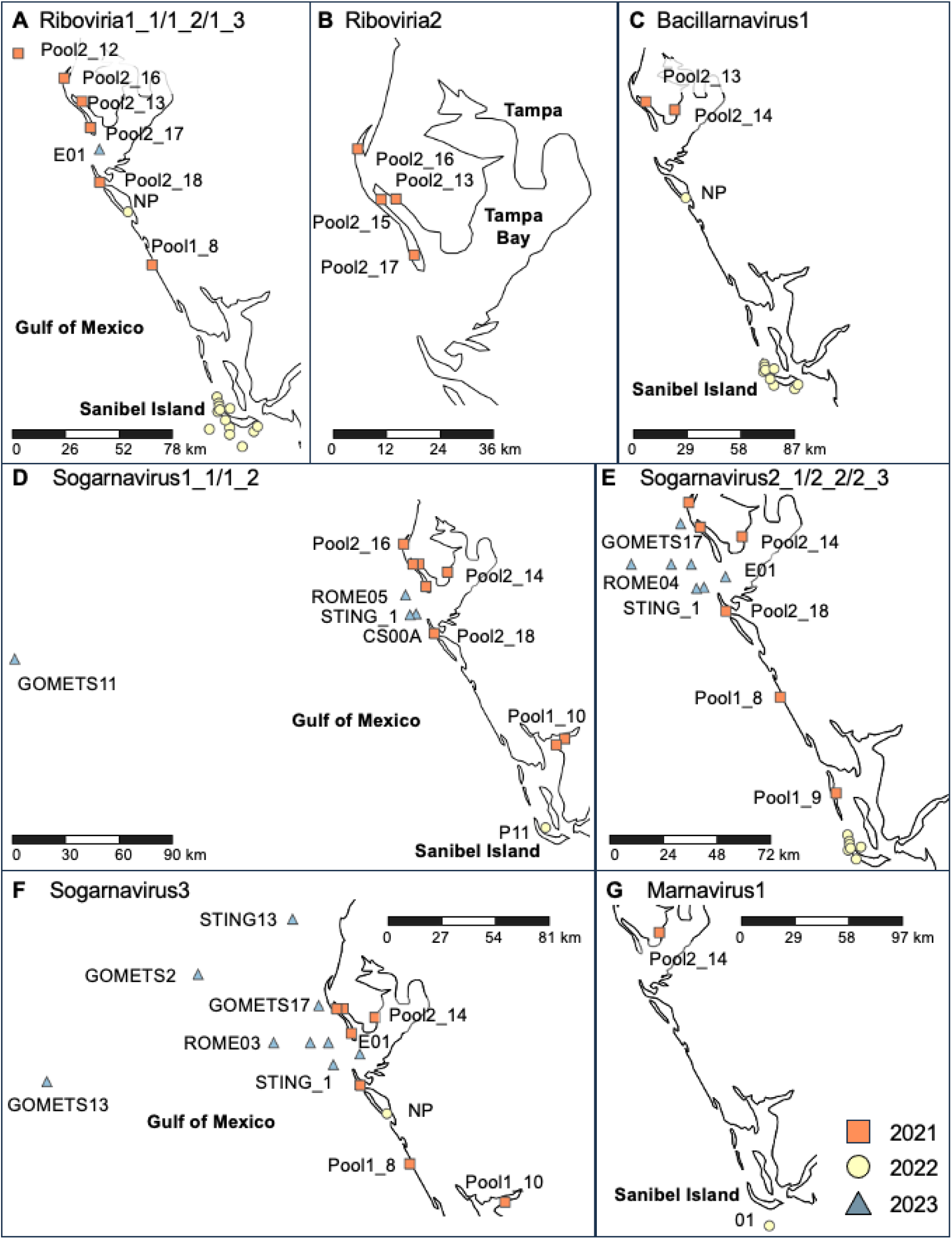
Map of locations where sequence fragments were amplified from the genomes (a) Riboviria1_1, Riboviria1_2, and Riboviria1_3; (b) Riboviria2; (c) Bacillarnavirus1; (d) Sogarnavirus1_1 and Sogarnavirus1_2 (e) Sogarnavirus2_1, Sogarnavirus2_2, and Sogarnavirus2_3; (f) Sogarnavirus3; and (g) Marnavirus1. Marker shapes and colors correspond to the year of sample collection. Metadata for collected seawater samples are described in Table 1 and **Table S1**.

### Viral diversity

Alpha diversity, measured as the number of representative genomes from each taxon detected in each sample using RT-PCR, was significantly different between years, season, and month. Samples collected in 2021 during the prolonged *K. brevis* bloom event, and during the time of year when most blooms have subsided, had the highest alpha diversity, followed by samples collected in 2022 and then 2023, which represented the earlier and later parts of the 2022-2023 bloom event, respectively (**Figure 4a**). Alpha diversity was the highest in samples collected during the summer months, compared to samples collected during the winter and spring months (**Figure 4b-c**). Alpha diversity was also positively correlated with longitude (**Figure 4e**), but not with latitude and depth. Additionally, alpha diversity was significantly different across *K. brevis* cell count categories (**Figure 4d**), reflecting a positive correlation with *K. brevis* cell counts (**Figure 4f**). To further investigate relationships between these predictor variables, we performed multiple regression using alpha diversity as the response variable. The optimum model (adjusted R^2^=0.4945, *p*=0.00001538) contained two predictor variables, including month (*p*=0.002) and *K. brevis* cell count category (*p*=0.0348).

**Figure 4.**
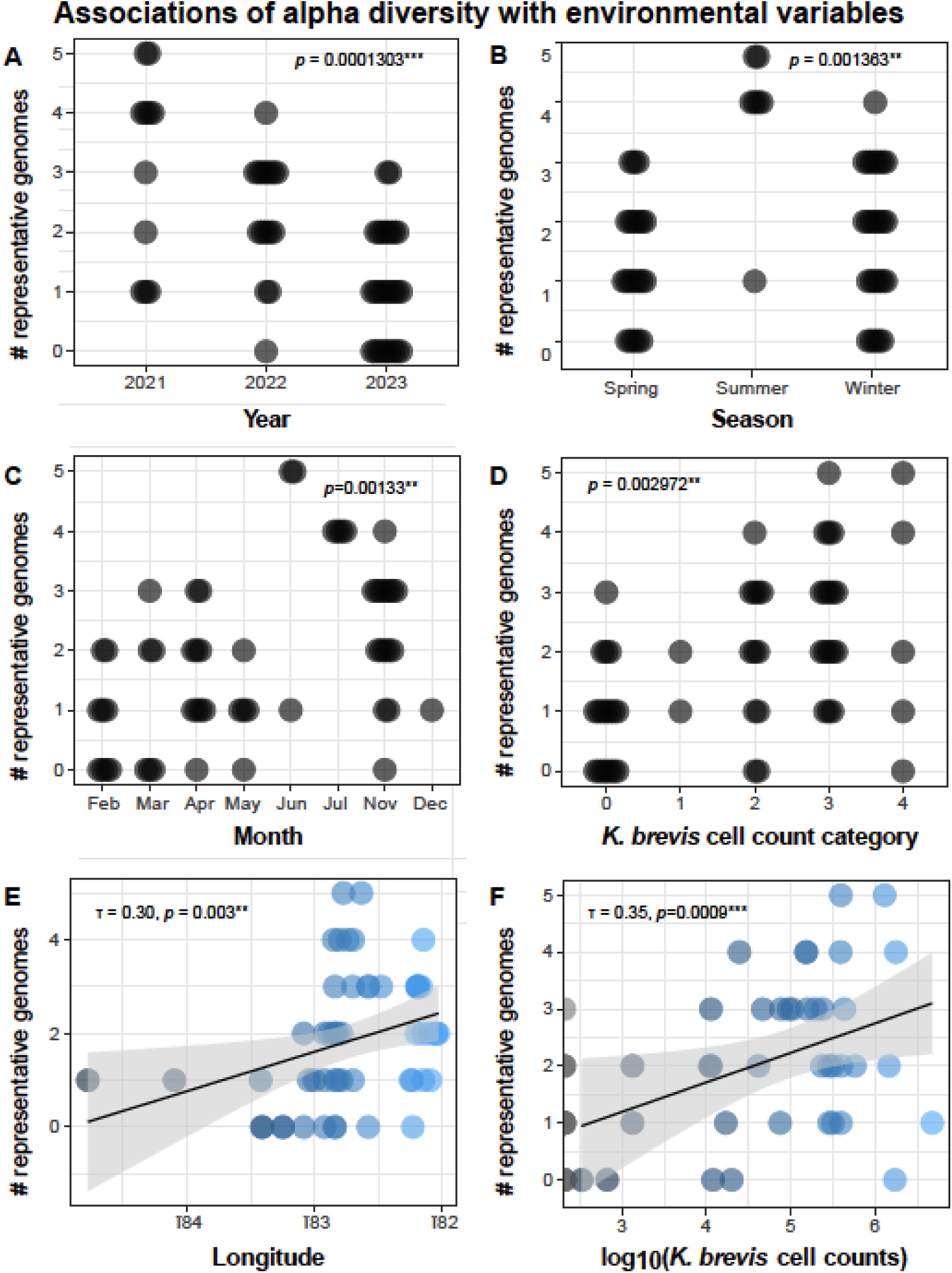
Statistical associations of alpha diversity with environmental variables, including (a) year; (b) season; (c) month; (d) *K. brevis* cell count category, (e) longitude, and (f) *K. brevis* cell counts (cells/L) represented as log_10_(*K. brevis* cell counts). Each marker represents a seawater sample and markers in (a-d) are arranged in beeswarm plots to reduce overlap. Marker colors correspond to the categories (a-d) or numerical values (e-f) of environmental variables. *p*-values in (a-d) are determined using Kruskal-Wallis tests, while Kendall’s τ correlation coefficient and *p*-values are reported in (e-f). ** denotes a *p*-value of ≤0.01 and *** denotes a p-value of ≤0.001.

Beta diversity was significantly correlated with latitude, longitude, year, season, month, and *K. brevis* cell count category (**Figure 5**). Multiple regression using *K. brevis* cell counts as the response variable generated an optimum model (adjusted R^2^=0.7887, *p*=8.035×10^−13^) with two predictor variables, including month (*p*=0.026) and the presence of sequences amplified from the genome Riboviria2 (*p*=0.039).

**Figure 5.**
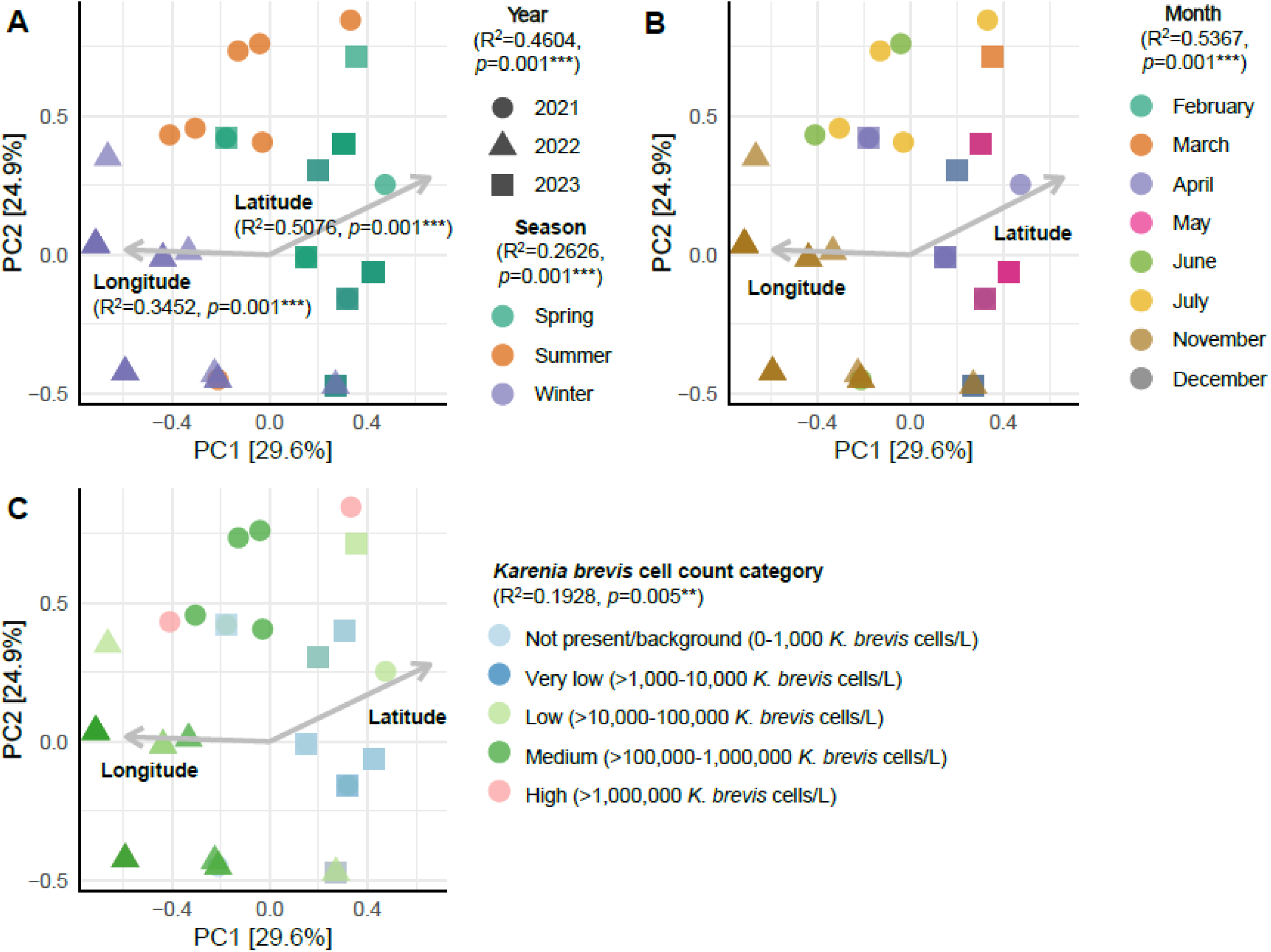
Principal components analysis (PCA) biplot showing the significant effects of longitude, latitude, (a) year and season; (b) month; and (c) *K. brevis* cell count category on the clustering of all field-collected samples. Each point in the PCA biplot represents a seawater sample and marker shapes correspond to the year of sample collection. Percentage values in square brackets on each axis represent the relative contribution (eigenvalue) of each axis to the total inertia (variation) in the dataset. The goodness-of-fit R^2^ value and *p*-value for each variable is shown on the biplots. ** denotes a *p*-value of ≤0.01 and *** denotes a *p*-value of ≤0.001.

To explore whether ssRNA viruses identified from field samples in this study could also be associated with cultivated *Karenia* species, we performed RT-PCR assays on *Karenia* spp. cell cultures routinely maintained in the laboratory at FWRI, including 17 *K. brevis* strains (CCFWC 242, 254, 257, 258, 261, 267, 121, 123, 124, 125, 126, 1010, 1012, 1013, 1014, 1016, and 1021), *K. mikimotoi* strain CCFWC 67, *K. papilionacea* strain CCFWC 1020, and *K. umbella* strain 1019 B2 (**Table S3**). These samples did not yield visible PCR products with any primer pair.

## Discussion

In this study, we recovered complete or near-complete genomes of ssRNA viruses from seawater collected around Southwest Florida during a long *K. brevis* bloom event that began in 2020 and lasted throughout most of 2021 (61). We developed targeted RT-PCR assays for these genomes to examine their occurrence and ecology in additional seawater samples collected in the region during a subsequent and shorter *K. brevis* bloom event between October 2022 and May 2023 (61).

Most of the assembled viral genomes belonged to the order *Picornavirales* and family *Marnaviridae* (45). Genomes assigned to the genus *Sogarnavirus* and *Bacillarnavirus* exhibit synteny and significant RdRp and capsid sequence similarity to genomes of viral species known to infect bloom-forming diatoms. These include *Chaetarnavirus 2* (CspRNAV02), which infects *Chaetoceros* sp., and *Rhizosolenia setigera* RNA virus 01 (RsRNAV01), which infects *Rhizosolenia setigera* (64). On the other hand, the genome assigned to the genus *Marnavirus* was related to *Heterosigma akashiwo RNA virus* (HaRNAV), which infects the HAB-forming raphidophyte species *Heterosigma akashiwo* (40). The prevalence of these ssRNA viruses, particularly *Sogarnavirus* sp., in our field-collected samples, is not surprising given that these phytoplankton genera are commonly observed in Southwest Florida (FWC HAB Monitoring Database). *Chaetoceros* sp., *R. setigera*, and *H. akashiwo* are globally distributed (65–67) and were previously identified in a 16S rRNA gene-based metabarcoding survey of seawater samples collected in Tampa Bay during the June 2018 *K. brevis* bloom (68). *Heterosigma akashiwo* has also been isolated from Tampa Bay (69, 70). *Chaetoceros* sp. and *H. akashiwo* are known HAB species that are associated with fish kills, especially at salmonid aquaculture net pens in Washington State (71, 72). Diatom species from the *Chaetoceros* and *Rhizosolenia* genera (including *R. setigera*) contain spines that could physically damage fish gills through clogging, mechanical irritation, puncturing, and/or secondary bacterial infections (67, 72). Although *R. setigera* is generally not considered a HAB species, there have been reports of fish mortalities caused by *R. setigera* blooms in Canada (73) and China (74). Algal viruses typically exhibit intraspecies host specificity with variations in strain specificity (75, 76). In contrast, an algal species can be infected by multiple strains of a virus, and strains of an algal species can differ in virus susceptibility (75, 76). Currently, the nature and impacts of phytoplankton-phytoplankton and phytoplankton-microbial interactions on *K. brevis* blooms in Florida are poorly understood. While phytoplankton, bacterial, and archaeal community composition was investigated during the June 2018 bloom (68), our study provides the first analysis on the diversity of RNA viruses associated with *K. brevis* blooms and suggests that, like blooms of *K. brevis*, associated viral assemblages are complex and dynamic.

In addition to genomes belonging to the *Marnaviridae* family, we also recovered genomes assigned to unclassified *Riboviria*. These genomes likely represent new viral taxa that are related to non-retroviral dinoflagellate-infecting +ssRNA virus (dinoRNAV) prevalent in coral holobionts (62, 77, 78) and distantly related to HcRNAV infecting the HAB-forming dinoflagellate species *Heterocapsa circularisquama* (63). Similar to dinoRNAV (62, 78) and HcRNAV (63), the unclassified *Riboviria* genomes contain two ORFs encoding RdRp and MCP. DinoRNAV has not been isolated in culture, but meta’omics analysis suggested that these viruses are endogenized within the genomes of coral dinoflagellate symbionts belonging to the family Symbiodiniaceae, particularly the genus *Symbiodinium* (63). The unclassified *Riboviria* genomes assembled in this study were most closely related to a dinoRNAV genome sequenced from a thermosensitive type C1 *Symbiodinium* culture (62). Furthermore, transcript expression of this dinoRNAV was downregulated under heat stress (62). In contrast, other dinoRNAV taxa were associated with heat-stressed coral samples (77) or showed higher MCP sequence diversity under heat stress (79). Because of their differential expression and diversity under thermal stress conditions, dinoRNAV are hypothesized to contribute to symbiont colonization and coral bleaching, which is partly driven by climate change (79). Although the presence of the genome Riboviria2 was a significant predictor of *K. brevis* cell counts, the small and imbalanced sample size in this study precluded statistical analysis of whether *K. brevis* cell counts could predict the presence (*n*=4) or absence (*n*=50) of sequences from the genome Riboviria2. Furthermore, the observation of Ribovaria2 only during summer months of 2021, outside the typical bloom season, is intriguing; further sampling can help evaluate whether Ribovaria2 occurs seasonally even if additional blooms do not persist through summer. We were also unable to determine the host of these unclassified *Riboviria* species. RT-PCR screening on *Karenia* spp. (*K. brevis*, *K. mikimotoi*, *K. papilionacea*, and *K. umbella*) cultures maintained in the laboratory over extended time periods did not amplify sequence fragments from these unclassified *Riboviria* genomes. Future culturing, imaging, and single-cell RNA sequencing experiments on fresh seawater samples containing *K. brevis* blooms are critical to isolate, characterize, and identify the host species of the unclassified *Riboviria* and other ssRNA viruses identified in this study. Since temperature data were not recorded for all our seawater samples, we were also unable to explore the effects of temperature on the prevalence or abundance of these unclassified *Riboviria* taxa and other viral taxa identified in this study beyond seasonal climatology (80). Notably, month was a significant predictor of both RNA viral diversity and *K. brevis* cell counts, and higher viral diversity was observed during the bloom event that persisted into the summer. Temporal effects could correspond to environmental parameters, including temperature, salinity, light availability, and nutrient concentrations, which have been known to influence the growth rate and physiology of *K. brevis* (81) and the seasonality of *K. brevis* blooms (82).

Overall, our study provided valuable information on the diversity of ssRNA viruses associated with *K. brevis* blooms and unveiled new *Riboviria*-dinoflagellate associations in the waters of Southwest Florida. Viruses play important roles in phytoplankton dynamics and cell death that can influence bloom dynamics (24). While viral interactions with other phytoplankton species are relatively well-studied (68, 83–86), viruses infecting *K. brevis* and co-occurring phytoplankton species during bloom events have been poorly investigated. Long-term surveillance of *K. brevis* blooms and their microbial and viral community composition, coupled with mechanistic studies, will help elucidate the roles of viruses in bloom ecology and their responses to environmental factors, including climate change, in this region.

## Acknowledgments

This work was supported by Florida Fish and Wildlife Commission (FWC) Cooperative Red Tide Research Program, award #20035. We acknowledge the contribution of Arianna Rodriguez in field work and sample collection.

